# Serotonin blocker Ketanserin reduces coral reef fish *Ctenochaetus striatus* aggressive behaviour during between-species social interactions

**DOI:** 10.1101/2023.01.20.524902

**Authors:** Virginie Staubli, Redouan Bshary, Zegni Triki

## Abstract

A multitude of species engages in social interactions not only with their conspecifics but also with other species. Such interspecific interactions can be either positive, like helping, or negative, like aggressive behaviour. However, the physiological mechanisms of these behaviours remain unclear. Here, we manipulated the serotonin system, a well-known neurohormone for regulating intraspecific aggressive behaviour, to investigate its role in interspecific aggression. We tested whether serotonin blockade affects the aggressive behaviour of a coral reef fish species (*Ctenochaetus striatus*) that engages in mutualistic interactions with another species, the cleaner fish (*Labroides dimidiatus*). Although this mutualistic cleaning relationship may appear positive, cleaner fish do not always cooperate and remove ectoparasites from the other coral reef fish (“clients”) but tend to cheat and bite the client’s protective layer of mucus. Client fish thus often apply control mechanisms, like chasing, to deter their cleaner fish partners from cheating. Our findings show that blocking serotonin receptors 5-HT2A and 5-HT2C with ketanserin reduced the client fish’s aggressive behaviour towards cleaner fish, but in the context where the latter did not cheat. These results are evidence of the involvement of serotonin in regulating aggressive behaviour at the between-species social interactions level. Yet, the direction of effect we found here is the opposite of previous findings using a similar experimental set-up and ecological context but with a different client fish species (*Scolopsis bilineatus*). Together, it suggests that serotonin’s role in aggressive behaviour is complex, and at least in this mutualistic ecological context, the function is species-dependent. This warrants, to some extent, careful interpretations from single-species studies looking into the physiological mechanisms of social behaviour.

## Introduction

A behaviour is usually termed ‘social’ if it affects not only the actor’s fitness but also the fitness of one or several recipients (Hamilton, 1964). Social behaviours can have both positive effects, such as grooming and helping, and negative effects, such as competition and aggression (Tinbergen, 2012). The functional definition of social behaviour implies that recipients can be conspecifics as well as heterospecifics (Oliveira and Bshary, 2021). However, such an inclusive approach may be challenged on a mechanistic level unless it can be shown that intra- and interspecific interactions are governed by the same underlying mechanisms (Oliveira and Bshary, 2021). Here, our focus is on physiological mechanisms such as endocrine systems. For instance, a well-studied social behaviour and its underlying endocrine mechanisms is aggression. Aggression can be either defensive or offensive, and it can serve to deter a predator, competing over limited resources (e.g., territory, food, shelter, and mates) and punishment-like behaviour to promote cooperation and prevent partners from defecting (Barnard, 2004; Raihani et al., 2012). Therefore, given the wide range of contexts wherein aggression may occur, most chemical messengers acting on the brain can be involved in regulating this behaviour.

Some chemical messengers are well studied in-depth than others for their role in the decision-making of aggression in a multitude of species and across taxa, such as steroid hormones, neuropeptides and neurotransmitters (Adkins-Regan, 2005; Reeder and Kramer, 2005; Ricklefs and Wikelski, 2002; Wingfield et al., 1998). In fact, several of these endocrine systems are phylogenetically ancient and well-preserved across taxa. One of the most ancient systems is that of the serotonin (Azmitia, 1999). Serotonin, or 5-hydroxytryptamine (5-HT), is a monoamine neurotransmitter that acts as a messenger among nerve cells. This neurotransmitter is also considered a hormone and can act through a family of receptors that can have either an inhibitory or excitatory role when activated. Decades of research on this molecule made it widely known for its imminent role in modulating aggressive behaviour in various species and taxa, like primates, including humans (Kuepper et al., 2010; Larke et al., 2016), reptiles (Larson and Summers, 2001; Summers et al., 2005), birds (Fachinelli et al., 1989; Sperry et al., 2003), fish (Stettler et al., 2021; Weinberger II and Klaper, 2014), and insects (Rillich and Stevenson, 2019). Nevertheless, how serotonin modulates aggression and whether it inhibits or enhances this behaviour still needs to be answered, given the system’s complexity and the broad contexts wherein aggression may occur (Olivier, 2004). Thus, case-by-case experimental studies using excitatory or inhibitory manipulations for targeted serotonin receptors and in well-defined ecological contexts can be a promising approach to unravelling the effects of serotonin on aggressive behaviour.

Here, we manipulated serotonin in a between-species social interactions context, an ecological context often overlooked in animal studies on aggressive behaviour and its mechanisms. A suitable study system is that of the mutualistic cleaner fish *Labroides dimidiatus* and its various coral reef fish partners (“clients”) that engage in iterated cleaning interactions for parasite removal (Losey, 1972). This relationship between cleaner fish and their clients is mutually beneficial, wherein cleaner fish gain food (Grutter, 1999) and client fish get rid of their ectoparasites, eventually boosting their overall health status (Clague et al., 2011; Demairé et al., 2020; Ros et al., 2020; Triki et al., 2016). However, despite the positive aspect of the relationship, conflict may occur as cleaner fish prefer to feed on the client’s protective mucus (Grutter and Bshary, 2003). As a control mechanism to deter their cleaner fish partner from cheating and encourage them to cooperate, client fish have two possible strategies. They either chase the biting partner or switch to another one for their next inspection, causing the loss of a potential food patch for the biting cleaner fish (Bshary and Grutter, 2002; Grutter and Bshary, 2003). This between-species social relationship, where the client-cleaner fish interactions fluctuate between positive and negative, makes it ideal to ask whether serotonin is a potential mediator of aggressive behaviour and, if so, how it affects this behaviour.

At least 14 different serotonin receptor types are being identified in mammals. Fish, in contrast, have three types (5-HT1, 5-HT2 and 5-HT7) with three subtypes of 5-HT1 and two subtypes of 5-HT2, the 5-HT2A and 5-HT2C, that have been identified so far (Mager et al., 2012; Norton et al., 2008; Schneider et al., 2012). Given that the role of serotonin is strongly dependent on the receptor system (see review by Siever, 2008), we opted for the molecule ketanserin (KET), a potent selective antagonist of serotonin receptors in fish that targets 5-HT2A and 5-HT2C receptors (Whitaker et al., 2011). Our study builds on previous work by Triki et al. (2017), where the serotonergic system emerged as a key candidate for the regulation of interspecific aggression in comparison to vasotocin (fish homologue of vasopressin) and isotocin (fish homologue of oxytocin) – which are two neurohormones known for their role in regulating intraspecific social behaviour (Adkins-Regan, 2005). That study used client fish monocle bream (*Scolopsis bilineatus*) as the focal subjects. Here, we followed a similar protocol as Triki et al., but we used a different species of client fish, the bristletooth surgeonfish (*Ctenochaetus striatus*). This aimed at facilitating the direct comparison of the findings and seeing whether ketanserin acts the same way in the two species when exposed to a similar ecological context, which is crucial to help disentangle the joint and separate roles of serotonin in regulating aggressive behaviour in the marine cleaning mutualism context across species. The set-up consisted in using wild-caught cleaner fish and their *C. striatus* clients, manipulating ketanserin in the client fish and testing how our treatment affected their aggressive behaviour towards the cleaner fish.

Furthermore, to investigate whether our treatment may have indirectly impacted the overall quality of the cleaner fish cleaning services, we measured the duration of the cleaning interactions, the amount of tactile stimulations and the cheating rate. Tactile stimulations occur when cleaner fish touches the client fish with its pelvic and pectoral fins, which can reduce the client fish’s stress levels (Soares et al., 2011). Cheating events, on the other hand, can be quantified as client fish body jolts caused immediately after the cleaner fish touches the client fish with its mouth, which indicates that the cleaner fish inflicted pain by taking a mucus bite (Bshary and Grutter, 2002). Triki et al. (2017) found that the blockade of serotonin 5-HT2A/2C receptors with ketanserin in *S. bilineatus* made them more aggressive towards cleaner fish in the absence of cheating events, but did not affect the aggression rates when a cleaner fish indeed cheated nor the quality of the cleaning interactions. Assuming that 5-HT2A and 5-HT2C receptors may have well-conserved functionality across species and context, we expected similar outcomes while testing a different client fish species.

## Methods

### Field site and animals

We conducted the study between July and September 2018 at the Lizard Island Research Station (LIRS) on the Great Barrier Reef (14.678436° S, 145.448280° E). Using barrier nets and hand nets, we caught 20 adult client surgeonfish, *Ctenochaetus striatus*, and 20 adult cleaner fish, *Labroides dimidiatus*. We housed all caught client fish individually in aquaria of 38 × 38.5 × 60 cm (height x width x length) and provided them with a shelter in the form of a polyvinylchloride cylinder (20 cm diameter and 20 cm length) and continuously aerated seawater. Similarly, we housed the cleaner fish individually in aquaria of 37 × 38 × 45 cm with polyvinylchloride pipes as shelter (2 cm diameter and 13 cm length). All client fish came from the same reef site (Osprey), whereas ten cleaner fish were from the reef site Northern horseshoe and the other ten from Corner beach. Immediately after capture, we deparasitized *C. striatus* using a freshwater bath for three minutes (Jones and Grutter, 2005) followed by an anti-helminthic aerated bath of Praziquantel (ICN Biomedicals Inc., Aurora, OH, USA) (1:100,000) overnight. We also measured fish body mass where *C. striatus* weighed on average 157.18 ± 49.57 grams (mean ± SD), *L. dimidiatus* from Northern horseshoe weighed 3.69 ± 0.99 grams, and *L. dimidiatus* from Corner beach weighed 3.25 ± 0.89 grams. We fed daily the *C. striatus* with a mixture of fish flakes and mashed prawns smeared on a Plexiglass plate and *L. dimidiatus* with mashed prawns smeared on Plexiglas plates. Furthermore, we allowed all caught fish to acclimate for at least two weeks before starting the experiment. By the end of the experiment, we returned and released all the surgeonfish *C. striatus* to their respective site of capture. The cleaner fish, *L. dimidiatus*, were used then in another study by Triki et al. (2020).

### Neuromodulator treatment

Based on the methods by Triki et al. (2017), we prepared the ketanserin treatment by dissolving 20 mg ketanserin (+)-tartrate (S006, Sigma-Aldrich) in 5 mL solution of 95% saline and 5% ethanol. We used the exact dosage as that by Triki et al. (2017), where we injected intramuscularly *C. striatus* with 10 μg/g of body weight. In the control condition, we injected the fish with a saline solution of identical volume to the ketanserin solution.

### Experimental set-up

We paired every focal client fish with a partner cleaner fish throughout the experiment on a size-based rule; for instance, the largest client fish formed a pair with the largest cleaner fish, and so forth. In a large round plastic tank measuring 105 cm in diameter and 42 cm high and containing continuously aerated seawater, we placed a pair of cleaner fish and client fish while keeping them separated by an opaque barrier (Supplementary Figure S1) and allowed them to acclimate to the experimental tank overnight. The focal fish received their treatment injection (saline or ketanserin) 10 min prior to the start of a trial. The trial started when the experimenter lifted the opaque barrier and allowed the two fishes to interact. We video-recorded these interactions for 15 min. At the end of the trial, both fish were returned to their respective home aquaria.

We ran the experiment for five consecutive days and tested eight cleaner-client pairs per day. Given that our experimental set-up is a matched design, we tested every cleaner-client pair twice, once with the ketanserin treatment and once with the control (saline), in a counterbalanced manner with a time gap of two days between the two injections.

### Behavioural analyses

The treatment identity of the video recordings was concealed from the experimenter who encoded these videos by renaming the files with running numbers (#1, #2, and so forth) to avoid potential subconscious bias. The experimenter used the open software CowLog 3.0.2. to extract the following cleaner-client interaction behaviours: (a) the total duration of the cleaning interactions; (b) the total duration of tactile stimulations; (c) the number of client fish’s jolts; and (d) whether the client fish chased the biting cleaner fish after a body jolt, i.e., provoked aggression. Furthermore, there was also chasing behaviour outside the context of the client fish responding to a cheating instance. These aggressions also occurred during cleaning interactions, but we could confirm on the video that no prior mouth contact could have caused the chasing. Therefore, we categorised these chasings as (e) unprovoked aggression.

### Data analyses

We used the open-access software R version 4.2.1 (R Core Team, 2022) to run all statistical analyses and generate the figures. Overall, we ran five different statistical models to test whether serotonin manipulation can affect the between-species social interaction patterns of the coral reef fish *C. striatus*. The first model was a Linear Mixed Effects model (LMER), where we fitted the duration of cleaning interactions as the response variable, treatment (ketanserin vs saline) as a categorical predictor variable, and the client-cleaner fish pair identity as a random factor. This is because every client-cleaner fish pair was tested twice, once when the client fish was injected with ketanserin and once with saline. Given that cleaner fish were caught from two different reef sites, we also added the reef site identity as a random factor. The second model was also an LMER model, with the duration of tactile stimulation as the response variable, treatment as a predictor, and standardised and log-transformed duration of cleaning interactions as a co-variate, while client-cleaner pair and cleaner fish reef site identities as random factors. The third model investigating client fish body jolts is a Generalized Additive Model for Location Scale and Shape (GAMLSS) because GAMLSS models support a wide range of data distribution than other statistical models (Stasinopoulos and Rigby, 2008). For instance, our model contained data that followed negative binomial distribution but contained several zeros. Therefore, we fitted a GAMLSS model with zero-inflated negative binomial error distribution for the count of client fish jolts as the response variable, treatment as a predictor, standardised and log-transformed duration of cleaning interactions as a co-variate, while client-cleaner pairs and cleaner fish reef site identities as random factors. Fourth, we fitted a GAMLSS with beta-inflated error distribution (data range was [0, 1]) since our response variable in this model was the proportion of provoked aggressions calculated from total jolts. The predictor variable was treatment, and client-cleaner pair and cleaner fish reef site identities were random factors. The final model was also a GAMLSS, but with zero-inflated Poisson distribution, wherein we fitted the count of unprovoked aggressions as the response variable, treatment as the predictor, and standardised and log-transformed duration of cleaning interactions as a co-variate, while client fish and cleaner fish reef site identities as random factors. We ran model diagnostics for all five models, and all of them fitted their respective assumptions, such as normality of residuals and homogeneity of variance. Please refer to our step-by-step code provided along with the data via the shared link in the Data and Code accessibility statement for further details.

## Results

We tested N = 20 individual client fish (*Ctenochaetus striatus*), but two did not interact with their partner cleaner fish in the saline treatment, yielding 38 video recordings. Our data analyses of the cleaning interaction patterns showed that our neuromodulator treatment did neither affect the overall duration of cleaning interactions (N = 38 videos; estimate β [low – high 95% credible interval] = −14 [−90 – 62.1], p-value = 0.703, Fig. 1a), the amount of tactile stimulations received (N = 38 videos; β = 15.2 [−4.67 – 35.1], p = 0.125, Fig. 1b), nor the frequency of being cheated by their partner cleaner fish (N = 38 videos; β = −0.432 [−2.32 – 1.30], p = 0.479, Fig. 1c).

**Figure 1.**
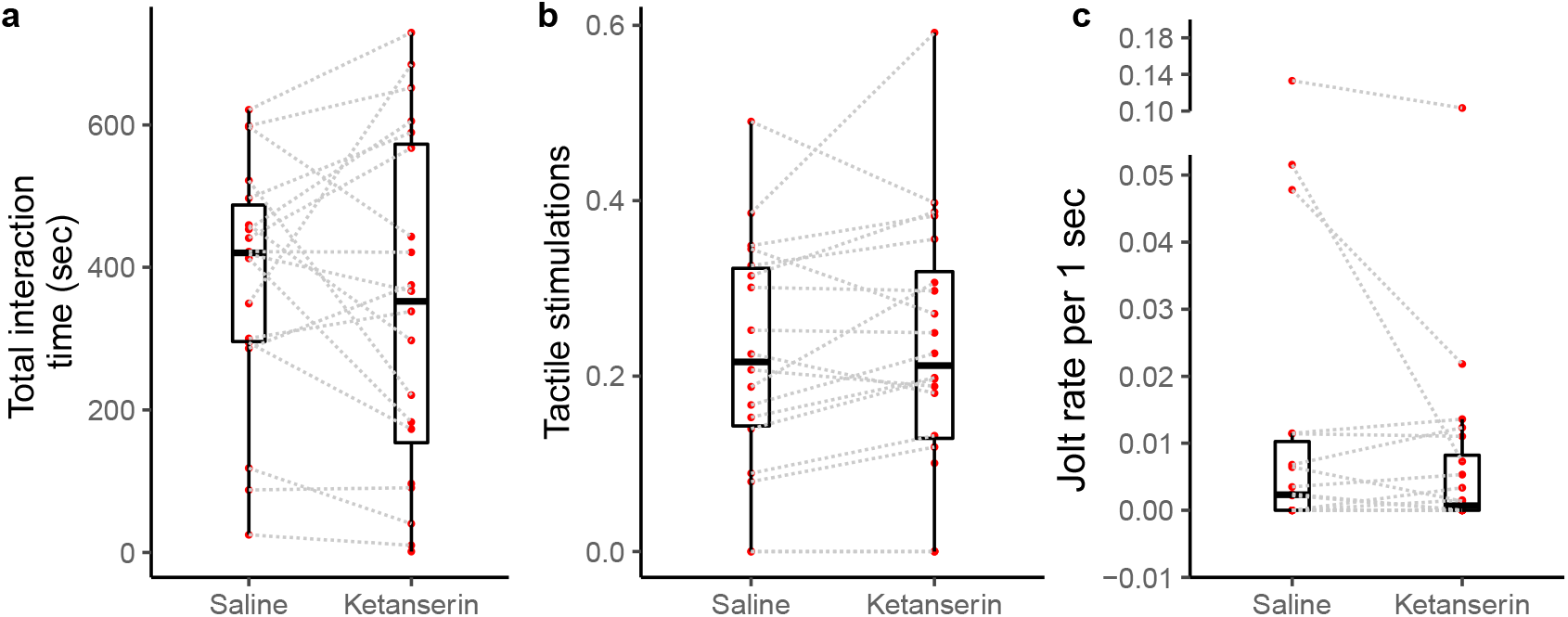
Cleaner-client interaction patterns as a function of neuromodulator treatment. Boxplots of median, interquartile and ranges of (a) total duration of the cleaning interactions per 15 min of observation; (b) the amount of tactile stimulation provided during these cleaning interactions, here, depicted as a time proportion measure for visualisation (see statistics for details about the analyses *per se*); and (c) the number of client fish body jolts occurring per 1 sec of interaction time (here as well we show the jolt measure as a rate for simplified visualisation (see statistics for details about the analyses *per se*). Also, the raw data points are depicted where every matched trial (individual client fish) is connected with a dashed line to visualise individual behavioural changes as a function of the neuromodulator treatment.

Notably, the client fish provoked aggression towards cleaner fish in the form of chasing the cleaner fish after a cheating event – often indicated by the client fish body jolt – was not affected by the neuromodulator treatment (N = 20 videos – as only 20 videos out of 38 had client fish showing at least one jolt event during the recording; β = −0.789 [−2.29 – 0.71], p = 0.318, Fig. 2a). Importantly, client fish aggressive behaviour in an unprovoked context, when they chase cleaner fish without the latter causing client fish to jolt, was significantly affected by the neuromodulator treatment (N = 38 videos; β = −0.774 [−1.30 – −0.248], p =0.006, Fig. 2b). The data showed that client fish receiving ketanserin injections were less likely to chase the cleaner fish compared to when they received saline injections, in the context where such chasing was unprovoked by the cleaner fish. Further statistical outcomes are available in the Electronic Supplementary Material (Table S1).

**Figure 2.**
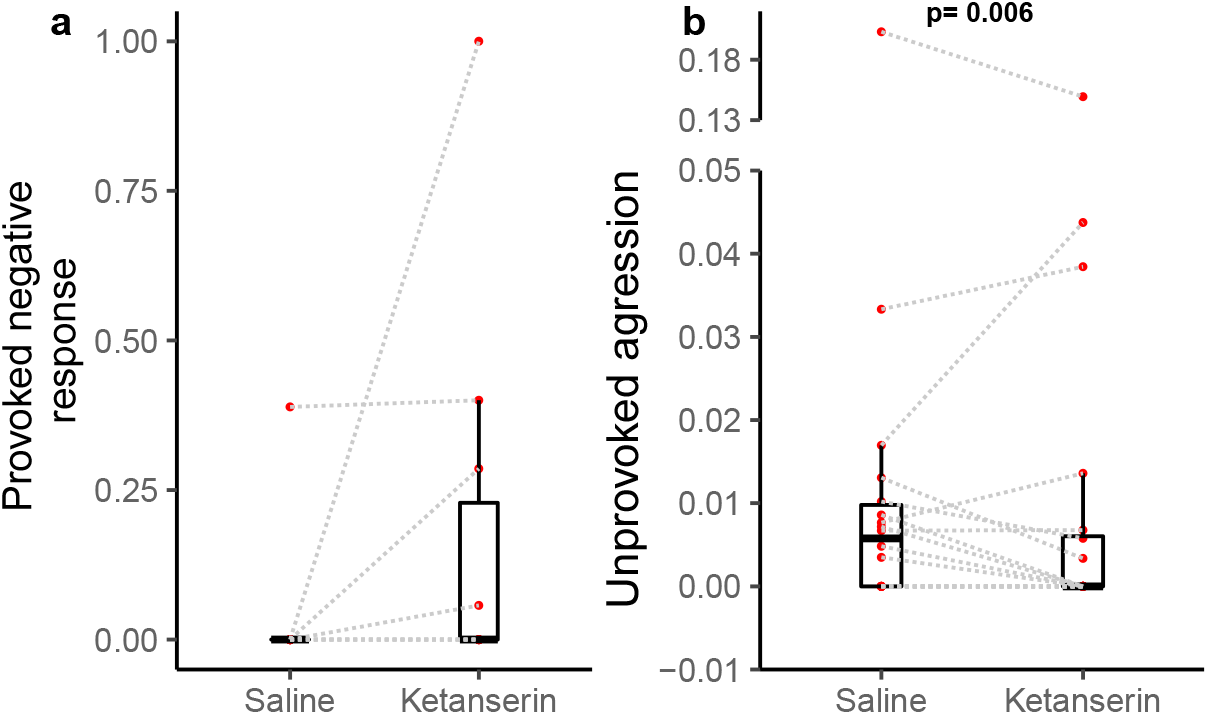
Client fish’s aggressive behaviour towards the cleaner fish as a function of neuromodulator treatment. Boxplots of median, interquartile and ranges of (a) provoked aggression, which is the proportion of client fish chasing their cleaner fish partner after a body jolt; and (b) unprovoked aggression occurring per 1 sec of interaction time (here we show the unprovoked aggression measure as a rate for simplified visualisation (see statistics for details about the analyses *per se*). Also, the raw data points are depicted where every matched trial (individual client fish) is connected with a dashed line to visualise individual behavioural changes as a function of the neuromodulator treatment.

## Discussion

We aimed to test whether the selective blockade of serotonin receptors, the 5-HT2A and 5-HT2C, affect coral reef fish aggressive behaviour in a between-species social interaction context. Although serotonin manipulation did not change the overall response of *C. striatus* to cheating events where most of their aggressive behaviour is expected to occur, it affected their “unprovoked” aggression, the rate by which they chased their cleaner fish partner without the latter having caused or provoked such attacks during the cleaning interactions. We found that the serotonin antagonist, ketanserin, reduced the rate of these unprovoked aggressive attacks, suggesting that in the client fish species *C. striatus*, serotonin action through the 5-HT2A/2C receptors reduces tolerance of non-conspecifics. Below, we discuss these findings in light of previous work that used a similar experimental set-up but with different client fish species. We also review some of the relevant literature to try to understand where our results fit in the big picture of the complex serotonin-aggression relationship.

In the study by Triki et al. (2017), they injected the client fish species *S. bilineatus* with ketanserin and compared their behaviour to those injected with saline as a control. In both Triki et al. (2017) and the current study, ketanserin did not affect the client-cleaner fish interaction quality or the client’s response to cheating. This direct comparison of two of the most common client reef fishes suggests that serotonin, via 5-HT2A/2C receptors, does not modulate key aspects, like client fish’s willingness to receive cleaning services or exerting punishment-like behaviours when their cleaner fish partner defects. In contrast, ketanserin increased the unprovoked aggression rate of *S. bilineatus* towards cleaner fish, while it decreased it in *C. striatus*. It is, however, noteworthy that in both studies, the jolt rate as an indicator of cheating was relatively low – here only half of the video observations showed at least one jolt rate. Cleaner fish in this experimental set-up tended to cooperate and rarely bit their clients. Having limited data to test how serotonin manipulation affects client fish response to cheating warrants careful interpretation. It is unclear whether serotonin does not regulate this behaviour in particular or we cannot see an effect due to insufficient data.

The absence of changes in the cleaning service quality, especially that of tactile stimulations, raises an interesting point, even if it was not necessarily the focus of this study. Biting cleaner fish often provide immediate tactile stimulations after a mucus bite as a reconciliation strategy (Bshary and Würth, 2001). Here, we did not see an evident increase in such behaviour in the control treatment, where they experienced significantly more unprovoked attacks compared to the ketanserin treatment. Although inconclusive, this finding points out the possibility that cleaner fish might not respond to an unjust punishment with an improved cleaning service.

For the unprovoked aggressive behaviour, our findings in reference to the study by Triki et al. (2017) show evident opposite effects of serotonin manipulation. Although unaware of the exact mechanisms yielding such results, we suggest the following potential explanations. For example, serotonin’s role in aggressive behaviour appears to be species-dependent. Although the system is phylogenetically ancient and well-preserved across invertebrate and vertebrate species, its function may have become adapted to the species. If the increase of serotonin decreases aggression in species A, for instance, it might have adopted an opposite role in species B. Examples from the literature support this point. For example, in rainbow trout, an increase in serotonin through dietary manipulation had an inhibitory effect on their aggressive response to an intruder (Winberg et al., 2001). Similarly, in several reptile species, increased serotonin levels in the brain are accompanied by enhanced aggression (Matter et al., 1998; Summers and Greenberg, 1995). In other species, like pigeons and song sparrows, increased serotonin yields lower aggression instead (Fachinelli et al., 1989; Sperry et al., 2003). Even within the same clade, we can see how serotonin mediates aggression in opposite directions. As such, in coral reef fish, the bluehead wrasse, *Thalassoma bifasciatum*, fluoxetine treatment – a selective serotonin reuptake inhibitor that yields higher serotonin activity – reduces chasing rates towards intruding conspecifics (Perreault et al., 2003), while in another coral reef fish species, the bicolour damselfish, *Pomacentrus partitus*, higher serotonin activity correlates positively with the number of aggressive attacks (Winberg et al., 1996). Thus, a logical explanation of our findings is that serotonin functionality in *C. striatus* differs from *S. bilineatus*. Therefore, it appears that serotonin’s role in regulating interspecific aggressive behaviour is complex; hence, any results from single-species studies cannot be generalised.

Following the blockade of the 5-HT2A/2C receptors, serotonin might have decreased *C. striatus* chasing attacks by becoming more available at the synaptic level to bind with and activate other sets of receptors like the 5-HT1A. This receptor is well-known for its role in reducing intraspecific aggression, especially in fish, like bluehead wrasse, *T. bifasciatum*, the fighting fish *Betta splendens*, and even in the cleaner fish *L. dimidiatus*, where excitation or inhibition of this receptor decreases and increases aggressive behaviour towards conspecifics, respectively (Clotfelter et al., 2007; Paula et al., 2015; Perreault et al., 2003). Moreover, we know that serotonin receptors can vary in their amount and distribution throughout the central nervous system, (Olivier, 2004). It is thus possible that quantitative variability in serotonin receptors between *S. bilineatus* and *C. striatus* drove the documented differences. Only extreme measures, like invasive methods involving killing the test subjects and performing immunoreactivity analyses (Frankenhuis-van den Heuvel and Nieuwenhuys, 1984), would show how serotonin agonists and antagonists affect their target receptors as well as the indirect effects on the other non-target serotonin receptors.

Finally, what adds to the complex role of serotonin in aggressive behaviour is the dose-dependent effect. Serotonin, among other neuromodulators, has an inverted U-shape function (Cano-Colino et al., 2014; Olivier, 2004; Stettler et al., 2021). Therefore, the increase or decrease in these neuromodulators can yield similar behavioural outputs if they are at the two ends of the inverted U-shaped curve. *S. bilineatus* and *C. striatus* might have responded differently to the same dose of ketanserin if their dose-dependent curve functions were not similar. Experimentally testing for this dose-dependent effect can help investigate species differences in serotonin function. However, this requires significant logistics that pose enormous practical challenges, especially for studies using wild animals.

Despite the opposing effects of ketanserin on the client fish’s unprovoked aggression towards cooperative cleaner fish, we conclude that serotonin does affect the client’s tolerance levels to proximity by cleaner fish, similar to what has been described in the literature about serotonin’s role in conspecifics’ tolerance and close contact (Insel and Winslow, 1998; Stettler et al., 2021). This suggests that the serotonin system has been co-opted to regulate aggression in general, both with individuals of the same or different species. The results thus contribute to the still relatively scarce evidence that interspecific interactions may be labelled as ‘social’ not only from a functional but also from a mechanistic perspective (Oliveira and Bshary, 2021).

## Supporting information

Electronic Supplementary Material

## Data availability

The data and code that support the present study’s findings are available in the repository Figshare (DOI: https://doi.org/10.6084/m9.figshare.11835690).

## Ethical note

The University of Queensland Animal Ethics Committee (AEC) approved the study under permit number: CA 2018/08/1222.

## Conflict of Interest

All authors declare that they have no conflict of interest.

## Acknowledgements

We thank the LIRS directors and staff for their support and friendship. Financial support was from the Swiss National Science Foundation (grant numbers 310030B_173334/1 to RB and PZ00P3_209020 to ZT).

## Notes

### Competing Interest Statement

The authors have declared no competing interest.

