## Supplementary material for "Serotonin blocker Ketanserin reduces coral reef fish *Ctenochaetus striatus* aggressive behaviour during between-species social interactions": Electronic Supplementary Material

**Author’s affiliation:**

### **^1^ Behavioral Ecology Laboratory, Faculty of Science, University of Neuchâtel, Emile-Argand 11, 2000 Neuchâtel, Switzerland**

^2^ Institute of Ecology and Evolution, University of Bern, Baltzerstrasse 1, 3012 Bern, Switzerland

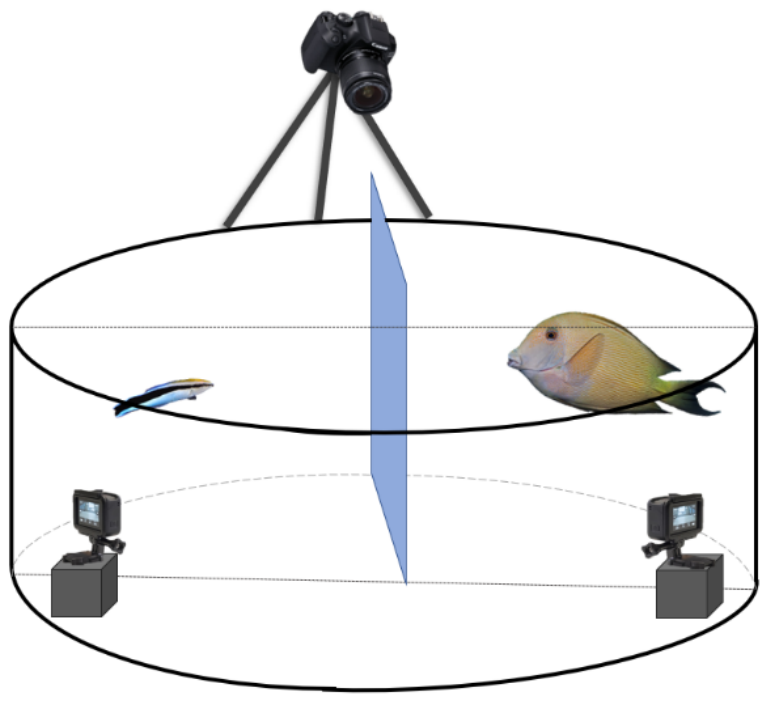

**Figure S1: Schematic representation of the experimental set-up to record the behavioural interactions between the client and cleaner fish.** Every client-cleaner fish pair was placed in a round plastic grey tank with a Plexiglas separation between the two fish to allow them to acclimate to the set-up. It is important to mention that fish did not have access to the two small GoPro® placed inside the tank. For that, we have placed a see-through barrier to prevent access. At the top, we installed a third camera, 5-Olympus®, to have a different view of the tank. Figure prepared by VS.

| **Statistical model class (distribution)** | **Measured behaviour (response variable)** | **Treatment (ketanserin vs saline)** | | | | **Covariate (Duration of cleaning interactions)** | | | **Explained variance (marginal and conditional R^2^ or pseudo-R^2^)** |
| --- | --- | --- | --- | --- | --- | --- | --- | --- | --- |
|  |  | **estimate** | **95 % Confidence level [low – high]** | | **p-value** | **estimate** | **95 % CI** | **p-value** |  |
| LMER (gaussian) | Duration of cleaning interactions | -14.0 | -90.0 – 62.1 | 0.703 | | - | - | - | 0.01 – 0.77 |
| LMER (gaussian) | Duration of tactile stimulations | 15.2 | -4.67 – 35.1 | 0.125 | | **52.1** | **30.1 – 74.0** | **< 0.001** | **0.41 – 0.88** |
| GAMLSS (Zero Inflated Negative Binomial) | Number of jolts | -0.432 | -2.32 – 1.30 | 0.479 | | 1.45 | 4.49 – -1.59 | 0.357 | 0.20 |
| GAMLSS (Beta inflated distribution ) | Provoked aggression (*chasing the cleaner fish after a cheating event*) | -0.789 | -2.29 – 0.71 | 0.318 | | - | - | - | 0.04 |
| GAMLSS (Zero Inflated Poisson) | Unprovoked aggression (*aggression towards cleaner fish without not preceded by a cheating event*) | **-0.774** | **-1.30 – -0.248** | **0.006** | | **-0.867** | **-1.27 -0.467** | **< 0.001** | **0.36** |

**Table S1.** Summary table with statistical outcomes from the different models testing changes in client fish behaviour as a function of the hormonal treatment manipulation where N = 20 client fish were injected with ketanserin and saline (as a control) in a matched design. The statistical significance level was set at alpha ≤ 0.05. Model name abbreviations are LMER = linear mixed effects model; and GAMLSS = Generalized Additive Models for Location Scale and Shape. Please refer to the main text for a detailed description of the measured behaviours.
